# Exercise serum promotes DNA damage repair and upregulates DNA repair gene *PNKP* in colon cancer cells

**DOI:** 10.1101/2025.06.26.659251

**Authors:** Samuel T Orange, Emily Dodd, Sharanya Nath, Hannah Bowden, Alastair Jordan, Hannah Smith, Ann Hedley, Ifeoma Chukwuma, Ian Hickson, Sweta Sharma Saha

**Author notes:** **Corresponding author** Dr Samuel T. Orange, School of Biomedical, Nutritional and Sport Sciences, Faculty of Medical Sciences, Newcastle University, Newcastle upon Tyne, NE2 4DR, UK, United Kingdom, **Email:**.

## Abstract

Exercise protects against colon cancer progression, but the underlying biological mechanisms remain incompletely understood. One proposed mechanism is the release of bioactive molecules into the interstitium and systemic circulation during exercise, which may act directly on precancerous or tumour cells to suppress DNA damage, inhibit proliferation, and preserve genomic stability. Here, we evaluated the effects of exercise-conditioned human serum on DNA damage kinetics and transcriptomic signatures in colon cancer cells. Blood samples were collected from 30 participants (age 50–78 years, body mass index ≥25 kg/m^2^) before and immediately after a maximal incremental cycling test. LoVo cells were exposed to pre- or post-exercise serum, treated with 2 Gy irradiation, and assessed for γ-H2AX foci over 24 hours. Compared to pre-exercise serum, post-exercise serum significantly reduced γ-H2AX foci at 6 hours (*p*=0.024) and decreased the area under the curve (AUC, *p*=0.014), indicating accelerated DNA repair. Post-exercise serum also increased expression of the DNA repair protein *PNKP* in LoVo cells, both with and without irradiation (*p*=0.007 and *p*=0.029, respectively). Transcriptomic analysis revealed upregulation of mitochondrial energy metabolism and downregulation of cell cycle and proteasome-related pathways. These findings suggest that acute exercise elicits systemic responses that enhance DNA repair and shift colon cancer cells towards a less proliferative transcriptomic state under sublethal genotoxic stress, offering a potential mechanistic explanation for the protective effects of exercise against colorectal carcinogenesis.

## 1 INTRODUCTION

Regular exercise suppresses colon cancer progression.^1–3^ Supervised exercise following adjuvant chemotherapy improves disease-free survival,^4^ and voluntary exercise reduces chemically-induced intestinal tumours in preclinical models.^1,2^ Putative biological mechanisms include enhanced immunosurveillance and reduced chronic inflammation and insulin resistance^5^. However, targeting these pathways has shown limited efficacy in improving colon cancer outcomes,^6–8^ and both chronic inflammation and insulin resistance appear to be regulated primarily by adiposity rather than exercise *per se*.^9,10^ Exercise improves colon cancer outcomes without changing adiposity,^4^ suggesting that additional mechanisms may be involved.

Emerging evidence demonstrates that bioactive molecules (proteins, nucleic acids, metabolites) released into the systemic circulation during exercise can act directly on cancer cells to inhibit tumour progression.^5,11^ These exercise-induced molecules, known as ‘exerkines’, exert systemic effects through endocrine-like signalling. We previously showed that exposing colon cancer cells to human serum sampled immediately post-exercise reduced cell proliferation by 6% compared to control serum.^12^ This was accompanied by a 25% decrease in γ-H2AX expression, a marker of DNA double-strand breaks (DSBs). Acute exercise also elevated serum IL-6—an established exerkine—and recombinant IL-6 reduced cell proliferation and DNA damage in a dose-dependent manner, mimicking the effects of exercise.^12^

Given that cancer cell lines rapidly attain new genetic variants in culture that accelerate proliferation,^13^ exercise-induced reductions in DNA damage may limit colon cancer cell proliferation by promoting genomic stability.^12^ This aligns with the oncogene-induced DNA damage model for cancer development, which proposes that persistent DNA DSBs drive cancer progression by increasing the propensity of acquiring genetic mutations that enhance cell proliferation and metastatic potential.^14^

Defining how exercise influences DNA damage repair in colon cancer may yield insights into its protective effects. A deeper mechanistic understanding could help identify biomarkers for risk stratification, inform exercise-mimetic drug development, or establish surrogate endpoints for trials. We therefore investigated the effects of exercise-conditioned human serum on DNA damage kinetics in irradiated colon cancer cells and examined associated transcriptomic changes to explore potential mechanisms.

## 2 METHODS

### 2.1 Study design

LoVo colon cancer cells were exposed to pre- or post-exercise human serum, treated with 2 Gray (Gy) irradiation to induce DNA DSBs, and assessed for nuclear γ-H2AX foci formation over 24 hours. Serum samples were generated from 30 participants who performed an acute bout of exercise, with blood samples collected pre- and immediately post-exercise. Prior to the exercise bout, participants fasted overnight (>10 hours), avoided structured exercise (≥48-hours), alcohol (≥24□hours) and caffeine (≥12□hours), and arrived hydrated. All experiments used individual (non-pooled) serum samples. The study was approved by the Newcastle University Faculty of Medical Sciences Research Ethics Committee (ref: 02219).

### 2.2 Participants

Thirty apparently-healthy adults aged 50-78 years completed the exercise trial. Inclusion criteria were age ≥50 years and body mass index (BMI) 25-39.9 kg/m^2^. Exclusion criteria were: (i) pre-existing cardiovascular, metabolic, or renal disease; (ii) fasting blood glucose ≥7 mmol/L, (iii) hypertension (≥160/≥90□mm□Hg); (iv) previous stroke or treatment for malignancy; (v) respiratory disease with peak expiratory flow <300□L/min; or (vi) any physical condition that could be exacerbated by exercise. Participant socio-demographics, medical history, BMI, waist circumference, body composition (Seca mBCA 515, Hamburg, Germany), peak expiratory flow, and fasting blood glucose were recorded (**Table 1)**.

**Table 1.** Participant characteristics.

|  | Total (n=30) | Male (n=18) | Female (n=12) |
| --- | --- | --- | --- |
| Age (years) | 64.1 ± 8.4 | 65.3 ± 7.1 | 64.3 ± 7.5 |
| Ethnicity |  |  |  |
| White British | 29 (97%) | 17 (94%) | 12 (100%) |
| Multiple ethnic groups | 1 (3%) | 1 (6%) | 0 (0%) |
| V̇O<sub>2peak</sub> (ml.kg<sup>-1</sup>.min<sup>-1</sup>) | 27.2 ± 6.7 | 29.7 ± 7.4 | 23.5 ± 2.9 |
| Body mass (kg) | 85.7 ± 11.0 | 90.3 ± 8.4 | 78.8 ± 11.0 |
| BMI (kg/m<sup>2</sup>) | 29.6 ± 2.9 | 29.5 ± 2.2 | 29.7 ± 3.8 |
| Waist circumference (cm) | 98.9 ± 8.8 | 102 ± 6.7 | 93.2 ± 9.3 |
| Body fat (%)<sup>a</sup> | 35.5 ± 6.9 | 30.5 ± 3.2 | 43.1 ± 2.2 |
| Fat-free mass (%)<sup>a</sup> | 64.5 ± 6.8 | 69.5 ± 3.2 | 57.1 ± 2.0 |
| Skeletal muscle mass (%)<sup>a</sup> | 29.3 ± 8.9 | 34.4 ± 2.3 | 22.6 ± 10.0 |
| Fasting BG (mmol/L) | 4.7 ± 0.8 | 4.7 ± 0.9 | 4.8 ± 0.7 |
| Peak expiratory flow (L/min) | 462 ± 120 | 536 ± 91.9 | 350 ± 50.8 |
| Data presented as mean ± SD or number (%) |  |  |  |

BG = blood glucose; BMI = body mass index; V̇O<sub>2peak</sub> = peak oxygen consumption.
n=15 due to bioelectrical impedance analysis not being available for all participants.

### 2.3 Sample size justification

Our primary outcome is difference in γH2AX foci after exposure to pre- vs. post-exercise serum. In our prior work, post-exercise serum reduced γ-H2AX expression in colon cancer cells by 25□±□47% (Cohen’s d_z_ = 0.53). A sample of n=30 participants provides 80% power to detect this effect with a two-tailed α = 0.05.

### 2.3 Exercise trial

After fasting overnight, participants performed a maximal incremental exercise test between 08:00-10:00 on an electronically braked cycle ergometer (VIAsprint 200P, Ergoline GmbH, Bitz, Germany). After a 3-minute warm-up against no resistance, work rate increased by 15-25 W·min^-1^ to reach volitional exhaustion in 10–12 minutes. Cadence was maintained at 60–75 rev·min^-1^; volitional exhaustion was defined as a ≥10 rev·min^-1^ drop in cadence for five consecutive seconds despite strong verbal encouragement. Breath-by-breath data (Vyntus CPX, Vyair Medical Inc., Chicago, IL, USA), heart rate (Polar H10, Polar Electro, Kempele, Finland), blood pressure, and rating of perceived exertion (RPE, 6-20 Borg scale) were recorded throughout. All but one participant achieved a peak respiratory exchange ratio (RER) of >1.10 (mean RER = 1.21 ± 0.10), confirming maximal effort.

### 2.4 Blood sampling

Venous blood samples (≈30□mL) were drawn from an antecubital vein before and within 1-minute of exercise cessation. Samples were collected in 10 mL Vacutainer serum tubes (BD, New Jersey), inverted 5-10 times, allowed to clot at room temperature for 30-minutes, centrifuged (1500g, 15-minutes), aliquoted (≈0.5 mL), and cryopreserved at −80°C.

### 2.5 Cell line

The LoVo cell line (LoVo, RRID:CVCL_0399) harbours *APC* and *KRAS* mutations but is wildtype for *TP53* and features the genomic instability phenotype microsatellite instability.^15^ Cells were purchased from Sigma-Aldrich (Dorset, UK) and cultured in DMEM/F12 (1:1) with 10% fetal calf serum, 1% glutamine, and 1% penicillin-streptomycin. For experiments, cells were passaged 4-8 times at ≈70-80% confluence and maintained at 37°C in a humidified atmosphere of 5% CO_2_. The cell line was authenticated using short-tandem repeat profiling (NorthGene, Newcastle upon Tyne, UK) and mycoplasma testing was performed every 3-months.

### 2.6 DNA damage repair

To assess DNA repair, nuclear γ-H2AX foci were quantified by immunofluorescence. LoVo cells were seeded at 5×10^4^ cells·well^−1^ in a 6-well plate for 48 hours, washed with phosphate-buffered saline (PBS), serum-starved for 2 h (0.2% bovine serum albumin [BSA] in DMEM), then treated with media containing: (i) 10% FCS; (ii) 10% pre-exercise serum; or (iii) 10% post-exercise serum. Paired pre- and post-exercise samples from the same participant (n=30) were used on the same 6-well plate at each timepoint post-irradiation to minimise interplate variability.^12^ After 1-hour serum stimulation, cells were irradiated with 2 Gy of x-ray radiation (0.8 minutes) and fixed at 5-minutes, 1-hour, 6-hours, and 24-hours. Cells were permeabilised with 0.2% Triton X-100 in PBS (PBS-Triton), blocked with PBS containing 2% BSA, 10% milk, and 10% goat serum, then incubated for one hour with mouse anti-phospho-H2AX antibody (1:1000, EDM Millipore) followed by Alexa Fluor 647-conjugated goat anti-mouse secondary antibody (1:1000, Abcam) in PBS-Triton with 2% BSA. Nuclei were counterstained with 4′,6-diamidino-2-phenylindole (DAPI, 1:1000). Images were captured using a Leica DM6 microscope (40× objective; LasX 3.4.2). γ-H2AX foci were quantified in ≥100 cells per coverslip using ImageJ (FIJI). Background counts from unirradiated controls were subtracted, and values were normalised to the 5-min post-irradiation timepoint. The area under the curve (AUC) for γ-H2AX over the 24-hours was calculated using the trapezoidal rule.

### 2.7 RNA sequencing

LoVo cells were treated with DMEM containing 10% pre- or post-exercise serum (n=10 and n=12 samples, respectively) for 6 hours, then harvested via trypsinisation, washed with PBS, and snap-frozen in liquid nitrogen. RNA was extracted and sequenced (Novogene, Cambridge, UK) using standard protocols. RNA extraction and sequencing were conducted by Novogene (Cambridge, UK). RNA quality was assessed using Nanodrop, Qubit, and Bioanalyzer instruments. Libraries were sequenced on an Illumina NovaSeq 6000, generating ∼25 million paired-end reads/sample. Reads were aligned to the human genome (GRCh38), and transcript abundance was quantified. Differential expression analysis was performed in R using DESeq2 (version 1.38.3), with significance set at false discovery rate (FDR) <0.05 (Benjamini-Hochberg corrected). Gene Ontology (GO) enrichment analysis was conducted using DAVID (https://davidbioinformatics.nih.gov/). Principal component analysis (PCA) was conducted on the top differentially expressed genes.

### 2.8. Real-Time Quantitative PCR

RNA (1 µg) was reverse-transcribed using the High-Capacity cDNA Reverse Transcription Kit with RNase inhibitor (Thermo Fisher, Cat #4374966) in 20 µL reactions with random hexamers. The reaction was incubated at 25°C for 10-minutes, 37°C for 2-hours, and 85°C for enzyme inactivation. Paired pre- and post-exercise samples (±2 Gy irradiation) were prepared for 11 participants. Quantitative PCR (qPCR) was performed using the Bio-Rad CFX Maestro system with GAPDH as the reference gene. Gene expression was analysed using the ΔCt method across technical replicates. Primer sequences are provided in **Table S1**. ΔCt values were plotted in GraphPad Prism (v10.4.2) and analysed by one-way ANOVA with Tukey’s post hoc test for multiple comparisons.

### 2.9 Statistical analysis

Linear mixed-effects models were used to compare γ-H2AX AUC between pre- and post-exercise serum conditions, with condition as a fixed effect and random slopes for participants. Additional models were fit separately at different timepoints (1-hour, 6-hours, 24-hours). Normality of residuals was assessed by visual inspection of histograms and Q–Q plots. Statistical significance was defined as p□<0.05. Analyses were performed in R (v4.4.3).

## 3 RESULTS

### 3.1 Exercise serum PROMOTES DNA damage repair in colon cancer cells

To investigate whether exercise serum modulates DNA repair, LoVo cells were stimulated with medium containing 10% human serum collected before and immediately after exercise. Following 1-hour serum exposure, cells were treated with 2□Gy x-ray irradiation, and γ-H2AX foci kinetics were assessed over 24 hours. Compared to pre-exercise serum, post-exercise serum significantly reduced γ-H2AX foci AUC (**p=0.014**), indicating an exercise-induced acceleration of DNA damage repair (**Figure 1**). Exercise serum also reduced γ-H2AX foci at 6 hours post-irradiation (−16.8 percentage points, 95% CI: -30.9 to -2.6 percentage points, ***p*=0.024**). Exercise-induced reductions in γ-H2AX foci levels at 1 hour (–17.1 percentage points, 95% CI: –34.4 to 0.35 percentage points, p = 0.059) and 24 hours (–13.0 percentage points, 95% CI: –26.1 to 0.02 percentage points, p = 0.054) post-irradiation did not reach statistical significance.

**Figure 1.**
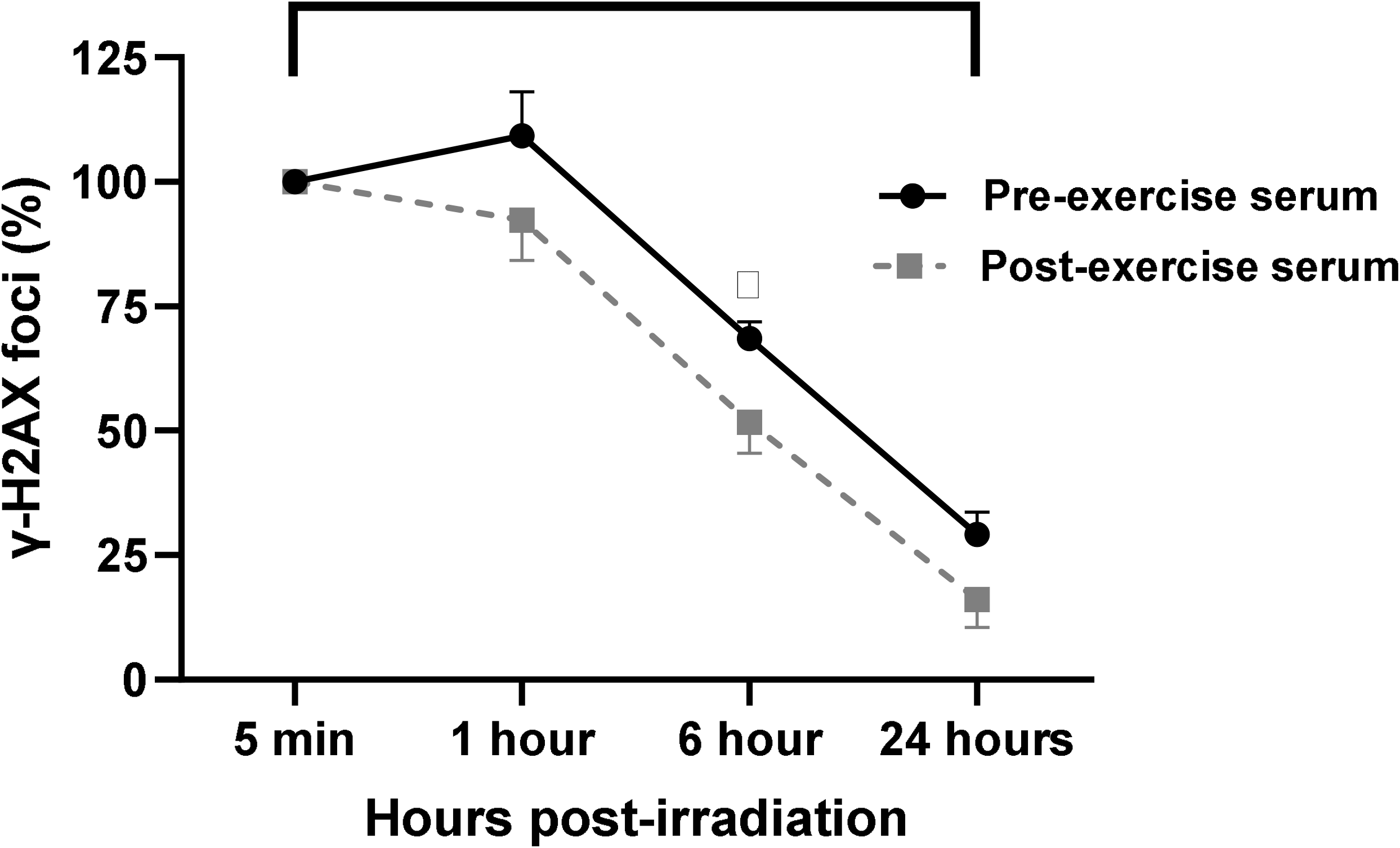
Exercise serum promoted DNA damage repair in irradiated colon cancer cells. LoVo cells were irradiated (2 Gy) following 1-hour stimulation with human serum collected before or immediately after acute exercise. γ-H2AX foci were quantified at 5 minutes, 1 hour, 6 hours, and 24 hours post-irradiation as a marker of DNA double-strand breaks. Data are expressed as a percentage of γ-H2AX foci relative to the 5-minute time point. Post-exercise serum significantly reduced γ-H2AX foci at 6 hours and decreased the area under the curve (AUC), indicating accelerated DNA damage repair. ^*^p < 0.05.

### 3.2 Exercise serum induces transcriptomic signatures of bioenergetic activation and proteostasis

To explore mechanisms underlying enhanced DNA repair, we performed RNA sequencing (RNA-seq) on LoVo cells stimulated with pre- or post-exercise serum. Post-exercise serum altered the expression of 1,364 genes compared to pre-exercise serum (FDR < 0.05), including 627 that were upregulated and 737 downregulated. Log□ fold changes (log□FC) ranged from –0.54 to 1.25 (median = – 0.11). GO analysis of upregulated genes revealed enrichment of biological processes related to protein synthesis (e.g., cytoplasmic translation, rRNA processing) and mitochondrial energy metabolism (e.g., oxidative phosphorylation, cellular respiration) (**Figure 2A**). Downregulated genes were enriched in pathways linked to cell cycle progression (e.g., cell division, G1/S transition of mitotic cell cycle) and proteasomal processes (e.g., protein polyubiquitination, deubiquitination) (Figure 2B). These transcriptional changes suggest that exercise serum promotes a cellular state of increased energy production and protein synthesis, alongside transient suppression of proliferation and protein degradation, which may reflect a transcriptional environment permissive of more efficient DNA repair. PCA of top differentially expressed genes confirmed clear separation between pre- and post-exercise conditions (**Figure 2B**).

**Figure 2.**
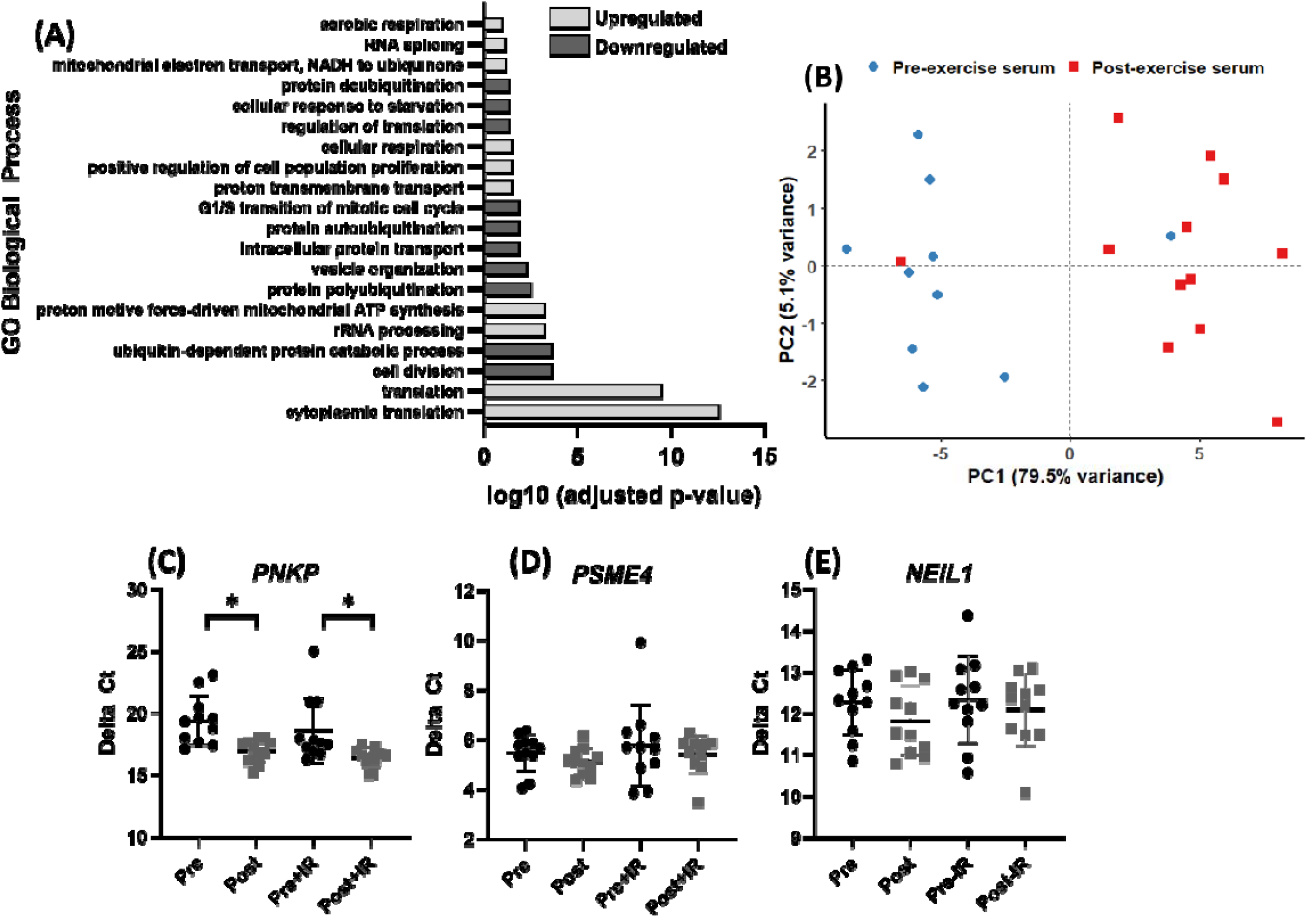
Exercise serum induces transcriptional signatures of enhanced bioenergetics and proteostasis and upregulates the DNA repair gene P*NKP* in colon cancer cells. **(A)** Gene Ontology (GO) enrichment analysis of genes upregulated in LoVo cells stimulated with post-exercise serum revealed over-representation of biological processes related to protein synthesis and mitochondrial energy metabolism and downregulation of cell cycle and proteasome-related pathways **(B)** Principal component analysis (PCA) of the top differentially expressed genes demonstrated clear separation between LoVo cells stimulated with pre- vs post-exercise serum, indicating distinct transcriptional profiles. **(C)** qPCR validation confirmed that post-exercise serum significantly upregulated *PNKP*, a key gene involved in DNA repair via base excision repair. **(D–E)** qPCR did not confirm differential expression of *PSME4* or *NEIL1*.

### 3.3 Exercise serum upregulates expression of DNA repair gene *PNKP*

To assess exercise effects on DNA repair gene expression, we selected *PNKP, PSME4*, and *NEIL1* for validation by qPCR based on transcriptomic changes and roles in DNA repair. *PNKP*, which encodes a DNA kinase-phosphatase essential for base excision repair, was significantly upregulated in the RNA-seq dataset (log□FC = 0.15, FDR = 0.035). qPCR confirmed increased *PNKP* expression following post-exercise serum stimulation: 1.9-fold (non-irradiated, ***p*=0.007**) and 4.5-fold (irradiated, ***p*=0.029**) relative to pre-exercise serum (**Figure 2C**). In contrast, the modest differential expression of *PSME4* (log□FC = –0.13) and *NEIL1* (log□FC = 1.07) observed in RNA-seq were not validated by qPCR (*p*>0.05, **Figures 2D–E**).

## 4 DISCUSSION

This study demonstrates that exercise-conditioned serum promotes DSB repair in colon cancer cells and induces transcriptomic signatures associated with enhanced bioenergetic activation and reduced cell cycle progression. These findings lend support to the hypothesis that systemic responses to acute exercise may suppress colon cancer progression, in part, by promoting DNA repair and shifting cells towards a less proliferative state under sublethal genotoxic stress.

Exposing colon cancer cells to post-exercise serum reduced γ-H2AX foci following treatment with 2 Gy ionising irradiation. This sublethal dose (2 Gy) induces DSBs and point mutations—including base substitutions, frameshifts, and small deletions— through misrepair mechanisms.^16,17^ Persistent sublethal genotoxic stress, particularly under conditions of error-prone repair, can promote the survival and expansion of genetically unstable clones.^14^ Enhancing DSB repair may therefore preserve genomic stability and limit clonal evolution. Although one proliferation-related pathway was upregulated, the overall transcriptional profile—marked by downregulation of cell cycle and proteasome-related genes—indicates a shift away from active proliferation, consistent with our prior findings.^18^ However, the relevance of enhanced repair in the context of cytotoxic therapies, which aim to induce irreparable DNA damage, remains uncertain and warrants further investigation.

Post-exercise serum upregulated mitochondrial pathways in LoVo cells, including oxidative phosphorylation (OXPHOS), electron transport, and aerobic respiration. Mitochondrial dysfunction contributes to colon cancer progression by increasing oxidative stress, impairing redox balance, and promoting genomic instability.^19^ The observed exercise-induced enrichment of mitochondrial metabolic pathways may suppress pro-tumour metabolic processes by enhancing OXPHOS and redox control, potentially reversing the glycolytic phenotype typical of many colorectal cancers.^20^ Indeed, enhancing mitochondrial biogenesis through AMPK-PGC-1α signalling—a pathway known to be activated by exercise^21^—can restore OXPHOS and inhibit colon cancer progression.^22^ Additionally, improved mitochondrial function may support more effective DNA repair, particularly energy-dependent single-strand break repair and non-homologous end joining (NHEJ), by maintaining cellular ATP levels and redox homeostasis essential for repair enzyme activity.

*PNKP* gene expression was upregulated in colon cancer cells stimulated with post-exercise serum, both with and without irradiation. *PNKP* encodes a critical enzyme involved in the repair of DNA strand breaks via base excision repair and NHEJ pathways. Although clinical transcriptomic data show inconsistent differences in *PNKP* expression between tumour grades,^23^ its function may be modulated by post-transcriptional regulation or the tumour microenvironment. In preclinical models, *PNKP* inhibition sensitises cells to lethal radiation and induces synthetic lethality in PTEN-deficient cancers.^24^ In contrast, the exercise-induced upregulation of *PNKP* observed here may reflect an adaptive response that supports DNA repair and genomic stability under sublethal genotoxic stress, such as that arising from oxidative or replicative damage within the tumour microenvironment or during early carcinogenesis. Further research should determine whether increased *PNKP* expression translates to functional effects on tumour behaviour *in vivo*.

Exercise is increasingly recognised as an adjunct therapy in cancer care. A recent trial found that a 3-year supervised exercise programme following chemotherapy improved disease-free survival in colon cancer, primarily through reduced recurrence and new tumours.^4^ These effects occurred without significant changes in body weight, suggesting that adiposity-related mechanisms are unlikely to explain the observed effects. Based on our findings, we hypothesise that regulation of DNA repair may be one mechanism through which exercise improves colon cancer outcomes.

We did not quantify serum analytes and therefore cannot identify the specific factors responsible for the exercise-induced reductions in DNA damage and transcriptional changes. Acute exercise triggers widespread changes in circulating proteins, metabolites, and nucleic acids that could influence DNA repair via metabolic, inflammatory, or regenerative pathways. However, as the serum metabolomic and proteomic responses to acute exercise are well characterised,^25^ further serum profiling was not pursued in this study.

In conclusion, exercise-conditioned serum enhanced DSB repair, upregulated *PNKP*, and induced transcriptional changes that reflect improved mitochondrial metabolism and reduced proliferation in colon cancer cells. These findings point to exercise-induced regulation of DNA repair as a potential mechanistic link between acute exercise and the suppression of colorectal carcinogenesis.

## Acknowledgments

We would like to thank Anna Turnbull, Aidan Perry, and Rosie Glossop for assisting with the exercise trials. We would also like thank the participants are taking part in the study.

## Author contributions

Samuel T. Orange: Conceptualisation, Funding Acquisition, Methodology, Formal Analysis, Investigation, Supervision, Project Administration, Writing—Original Draft.

Emily Dodd: Investigation, Data Curation, Writing—Review & Editing.

Sharanya Nath: Investigation, Data Curation, Writing—Review & Editing.

Hannah Bowden: Investigation, Data Curation, Writing—Review & Editing.

Alastair Jordan: Investigation, Data Curation, Resources, Writing—Review & Editing.

Anna Turnbull: Investigation, Writing—Review & Editing.

Hannah Smith: Investigation, Writing—Review & Editing.

Ann Hedley: Formal Analysis; Data Curation, Resources, Writing—Review & Editing.

Ifeoma Chukwuma: Investigation, Writing—Review & Editing.

Ian Hickson: Conceptualisation, Funding Acquisition, Methodology, Supervision, Writing—Review & Editing.

Sweta Sharma Saha: Conceptualisation, Funding Acquisition, Methodology, Investigation, Supervision, Project Administration, Validation, Writing—Review & Editing.

## Ethics statement

The study was approved by the Faculty of Medical Sciences Research Ethics Committee, part of Newcastle University’s Research Ethics Committee (ref: 02219). Written informed consent was obtained from all participants prior to data collection.

## Data availability statement

All data and code are available on the Open Science Framework project page (https://osf.io/2adyh/). Further information is available from the corresponding author upon request.

## REFERENCES

1. Ju J, Nolan B, Cheh M, et al. Voluntary exercise inhibits intestinal tumorigenesis in ApcMin/+ mice and azoxymethane/dextran sulfate sodium-treated mice. BMC Cancer. 2008;8:316. doi:10.1186/1471-2407-8-316

2. Kelly SA, Zhao L, Jung KC, et al. Prevention of tumorigenesis in mice by exercise is dependent on strain background and timing relative to carcinogen exposure. Sci Rep. 2017;7:43086. doi:10.1038/srep43086

3. Brown JC, Ma C, Shi Q, et al. Association between physical activity and the time course of cancer recurrence in stage III colon cancer. Br J Sports Med. 2023;57(15):965–971. doi:10.1136/bjsports-2022-106445

4. Courneya KS, Vardy JL, O’Callaghan CJ, et al. Structured Exercise after Adjuvant Chemotherapy for Colon Cancer. N Engl J Med. 0(0). doi:10.1056/NEJMoa2502760

5. Orange ST, Leslie J, Ross M, Mann DA, Wackerhage H. The exercise IL-6 enigma in cancer. Trends Endocrinol Metab. 2023;34(11):749–763. doi:10.1016/j.tem.2023.08.001

6. Jwk C, E S, Y D, et al. Aspirin after completion of standard adjuvant therapy for colorectal cancer (ASCOLT): an international, multicentre, phase 3, randomised, double-blind, placebo-controlled trial. Lancet Gastroenterol Hepatol. 2025;10(3). doi:10.1016/S2468-1253(24)00387-X

7. McNeil JJ, Gibbs *Peter, Orchard SG, et al. Effect of Aspirin on Cancer Incidence and Mortality in Older Adults. JNCI J Natl Cancer Inst. 2021;113(3):258–265. doi:10.1093/jnci/djaa114

8. Singh PP, Shi Q, Foster NR, et al. Relationship Between Metformin Use and Recurrence and Survival in Patients With Resected Stage III Colon Cancer Receiving Adjuvant Chemotherapy: Results From North Central Cancer Treatment Group N0147 (Alliance). The Oncologist. 2016;21(12):1509–1521. doi:10.1634/theoncologist.2016-0153

9. Brown JC, Compton SLE, Kang A, et al. Effects of exercise on inflammation, circulating tumor cells, and circulating tumor DNA in colorectal cancer. J Sport Health Sci. Published online March 17, 2025:101036. doi:10.1016/j.jshs.2025.101036

10. Kang DW, Lee J, Suh SH, Ligibel J, Courneya KS, Jeon JY. Effects of Exercise on Insulin, IGF Axis, Adipocytokines, and Inflammatory Markers in Breast Cancer Survivors: A Systematic Review and Meta-analysis. Cancer Epidemiol Biomark Prev Publ Am Assoc Cancer Res Cosponsored Am Soc Prev Oncol. 2017;26(3):355–365. doi:10.1158/1055-9965.EPI-16-0602

11. Orange ST, Jordan AR, Saxton JM. The serological responses to acute exercise in humans reduce cancer cell growth in vitro: A systematic review and meta-analysis. Physiol Rep. 2020;8(22):e14635. doi:10.14814/phy2.14635

12. Orange ST, Jordan AR, Odell A, et al. Acute aerobic exercise-conditioned serum reduces colon cancer cell proliferation in vitro through interleukin-6-induced regulation of DNA damage. Int J Cancer. 2022;151(2):265–274. doi:10.1002/ijc.33982

13. Ben-David U, Siranosian B, Ha G, et al. Genetic and transcriptional evolution alters cancer cell line drug response. Nature. 2018;560(7718):325–330. doi:10.1038/s41586-018-0409-3

14. Halazonetis TD, Gorgoulis VG, Bartek J. An oncogene-induced DNA damage model for cancer development. Science. 2008;319(5868):1352–1355. doi:10.1126/science.1140735

15. Berg KCG, Eide PW, Eilertsen IA, et al. Multi-omics of 34 colorectal cancer cell lines - a resource for biomedical studies. Mol Cancer. 2017;16(1):116. doi:10.1186/s12943-017-0691-y

16. Grosovsky AJ, de Boer JG, de Jong PJ, Drobetsky EA, Glickman BW. Base substitutions, frameshifts, and small deletions constitute ionizing radiation-induced point mutations in mammalian cells. Proc Natl Acad Sci. 1988;85(1):185–188. doi:10.1073/pnas.85.1.185

17. Yu Z qi, Zhang C, Lao X yuan, et al. [Long non-coding RNA influences radiosensitivity of colorectal carcinoma cell lines by regulating cyclin D1 expression]. Zhonghua Wei Chang Wai Ke Za Zhi Chin J Gastrointest Surg. 2012;15(3):288–291.

18. Orange ST, Jordan AR, Odell A, et al. Acute aerobic exercise-conditioned serum reduces colon cancer cell proliferation in vitro through interleukin-6-induced regulation of DNA damage. Int J Cancer. 2022;151(2):265–274. doi:10.1002/ijc.33982

19. Zhao Y, Guo X, Zhang L, Wang D, Li Y. Mitochondria: a crucial factor in the progression and drug resistance of colorectal cancer. Front Immunol. 2024;15:1512469. doi:10.3389/fimmu.2024.1512469

20. Qin R, Fan X, Huang Y, et al. Role of glucose metabolic reprogramming in colorectal cancer progression and drug resistance. Transl Oncol. 2024;50:102156. doi:10.1016/j.tranon.2024.102156

21. Sriwijitkamol A, Coletta DK, Wajcberg E, et al. Effect of Acute Exercise on AMPK Signaling in Skeletal Muscle of Subjects With Type 2 Diabetes. Diabetes. 2007;56(3):836–848. doi:10.2337/db06-1119

22. Hong R, Min S, Huang J, Zou M, Zhou D, Liang Y. High-dose vitamin C promotes mitochondrial biogenesis in HCT116 colorectal cancer cells by regulating the AMPK/PGC-1α signaling pathway. J Cancer Res Clin Oncol. 2025;151(5):167. doi:10.1007/s00432-025-06211-z

23. Tissue expression of PNKP - Summary - The Human Protein Atlas. Accessed June 11, 2025. https://www.proteinatlas.org/ENSG00000039650-PNKP/tissue

24. Sadat SMA, Paiva IM, Shire Z, et al. A synthetically lethal nanomedicine delivering novel inhibitors of polynucleotide kinase 3’-phosphatase (PNKP) for targeted therapy of PTEN-deficient colorectal cancer. J Control Release Off J Control Release Soc. 2021;334:335–352. doi:10.1016/j.jconrel.2021.04.034

25. Contrepois K, Wu S, Moneghetti KJ, et al. Molecular Choreography of Acute Exercise. Cell. 2020;181(5):1112-1130.e16. doi:10.1016/j.cell.2020.04.043

