## Supplementary material for "Exercise serum promotes DNA damage repair and upregulates DNA repair gene *PNKP* in colon cancer cells": Table S1

| **Table S1. List of qRT-PCR primers** | |
| --- | --- |
| **Target Gene Name** | **Primer Sequence** |
| PNKP_F | TCATGTATGGCTACAGGAAG |
| PNKP_R | AGGAGAAACAGCGTTTATTG |
| PSME4_F | TAGCTGTTTGTTTAACAGC |
| PSME_R | GGGCTTCTTGTATCTTTCAC |
| NEIL_F | AGCAGTTCAGGGAGAATG |
| NEIl_R | CAGATAGTTGCCAATGCC |
| GAPDH_F | TCGGAGTCAACGGATTTG |
| GAPDH_R | CAACAATATCCACTTTACCAGAG |
